# LOCAS: Multi-label mRNA *Loca*lization with Supervised Contrastive Learning

**DOI:** 10.1101/2024.09.24.614785

**Authors:** Abrar Rahman Abir, Md Toki Tahmid, M. Saifur Rahman

## Abstract

Traditional methods for mRNA subcellular localization often fail to account for multiple compartmentalization. Recent multi-label models have improved performance, but still face challenges in capturing complex localization patterns. We introduce LOCAS (Localization with Supervised Contrastive Learning), which integrates an RNA language model to generate initial embeddings, employs supervised contrastive learning (SCL) to identify distinct RNA clusters, and uses a multi-label classification head (ML-Decoder) with cross-attention for accurate predictions. Through extensive ablation studies and multi-label overlapping threshold tuning, LOCAS achieves state-of-the-art performance across all metrics, providing a robust solution for RNA localization tasks.

## 1 Introduction

The subcellular localization of messenger RNAs (mRNAs) is a critical process in the regulation of gene expression that ensures the spatial and temporal control necessary for proper cellular function [5]. mRNAs are not distributed uniformly within the cell but are instead localized to specific compartments. This localization allows for the precise control of protein synthesis, which is particularly important in complex cells. The asymmetric distribution of mRNAs has been shown to provide several advantages, including low transport costs and the prevention of ectopic protein activity during translocation [3, 9, 15].

The traditional methods used to study mRNA localization, such as in situ hybridization (ISH) and high-throughput RNA sequencing, have been invaluable in advancing our understanding of this process [1]. However, these methods are often time-consuming and costly, limiting their use in large-scale studies.

Initial computational approaches to mRNA localization framed the problem as a single-label classification task, where each mRNA was predicted to localize to only one specific compartment. RNATracker employed deep recurrent neural networks to predict mRNA localization [17]. iLoc-mRNA, utilized support vector machines (SVMs) to predict mRNA localization specifically in Homo sapiens, [22]. SubLocEP further refined predictions by concentrating on specific cellular compartments while remaining within the single-label classification framework [7]. However, they were inherently limited by the assumption that each mRNA localizes to only one compartment, which does not align with biological reality. Many mRNAs are known to localize in multiple compartments, fulfilling diverse roles within the cell [20, 7].

To address the limitations of single-label approaches, the field has shifted towards multi-label prediction models. Notable multi-label methods include DM3Loc, a deep learning-based model that utilized a multi-head self-attention mechanism to predict mRNA localization across multiple compartments simultaneously [15]. Moreover, Clarion [1] employed an ensemble learning strategy based on XGBoost, enhancing the accuracy and robustness of multi-label mRNA localization predictions by considering label correlations and utilizing advanced feature selection methods. Furthermore, Al-locator introduced the use of graph neural networks (GNNs) to incorporate RNA secondary structure information into the prediction model [6].

Multi-label data representation and prediction have recently been greatly studied in the field of computer vision and natural language processing. MultiSupCon [2] proposed an approach to understand the embedding space of sets of images with supervised contrastive learning with a projection embedding generated by an encoder. This helps the encoder to identify the distribution difference between different classes. In case of biological sequences, such initial embedding generation is constructured either with evolutionary [16], physiochemical [8, 14] features, or using a pretrained language model [11, 13] to generate embeddings. In this paper, we have proposed a similar approach for RNA sequence representation learning.

**Firstly**, we propose the integration of an RNA language model to generate an embedding space for RNA sequences in the RNA sub-cellular localization task. **Secondly**, we employ an effective supervised contrastive learning (SCL) algorithm to identify distinct clusters of RNA sequences, addressing the natural label overlap in this multi-label classification task using an overlapping threshold-based similarity score to identify similar clusters during SCL training. **Finally**, instead of using a simple fixed prediction head, we utilize a specialized classification head designed for multi-label tasks (ML-Decoder) that incorporates a cross-attention mechanism for classification. With these components, we propose multi-label mRNA **Loca**lization with **S**upervised Contrastive Learning (LOCAS). Through extensive ablation studies and hyper-parameter tuning during SCL training, we determine the optimal parameters for training LOCAS, achieving state-of-the-art performance across all metrics in RNA sub-cellular localization compared to previously reported approaches.

## 2 Methods

In this section, we first describe the datasets that are used for the RNA subcelluler localization task. Then we dive into the architectural details of different components: feature representation, encoder network, supervised contrastive learning for RNA sequences, prediction head and overall training pipeline.

### 2.1 Dataset Description

For the RNA subcelluler localization task we used the dataset from RNALocate [19] which is also used in DM3Loc [15]. The dataset comprises 17,298 unique mRNA sequences with six celluler locatations: Nucleus, Exosome, Cytosol, Ribosome, Membrane, and Endoplasmic Reticulum. However, each RNA sequence can belong to these multiple classes. We perform 5 fold cross-validation following [19, 15].

### 2.2 Initial Feature Representation and Encoder Network

Each RNA sequence in the dataset is first encoded with RNA language model RiNALMo [11]. For a given input to the RiNALMo, the output from the language model encoder is a sequence of vectors *F* = {[*CLS*], *f*_1_, *f*_2_, *f*_3_, …, *f*_*N*_ [*END*]}where each *f*_*i*_ *∈* ℝ^*d*^, with *d* = 1024. To get a sequence level representation of the RNA, we consider the [CLS] token of the transformer’s output (which is the first token of the output).

#### Encoder Network for Feature Space Transformation

The initial feature representation vector *F* is passed through an encoder network, *Enc*(·), that maps *x* to a representation vector 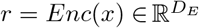, where *D*_*E*_ = 2048.

### 2.3 Supervised multi-label Contrastive Training

With the output from the encoder network *r* = *Enc*(*x*), we pass this through a projection layer 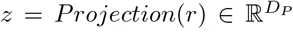, where *D*_*P*_ = 128. This projection layer is used for a contrastive training in the RNA dataset.

#### Self Supervised Vs Supervised Contrastive Learning

In a self-supervised setting, two distinct views of each RNA sequence are generated through augmentations like noise addition or random cropping, forming positive pairs, while other sequences in the batch serve as negative pairs. The challenge lies in the difficulty of reliably augmenting RNA sequences without compromising their structural integrity, unlike in natural language or vision domains. We propose a supervised contrastive learning approach using RNA subcellular localization labels. Similar sequences with the same label are considered positive pairs, while others are negative.

#### Overlapping Labels in multi-label Classification

In multi-label problems, such as RNA subcellular localization, the supervised contrastive loss has a notable limitation. Specifically, it only treats two sets of labels as belonging to the same class when they are an exact match. However, in multi-label scenarios, it is rare to encounter identical label sets within a single batch or even across the entire dataset. In subcellular localization task, where the possible labels are: Nucleolus, Exosome, Cytosol, Ribosome, Membrane, and Endoplasmic Reticulum (ER), each RNA sequence can have a combination of these labels.

To measure the degree of labels overlap between two sequences, we use the jaccard index, inspired from [10]. If two sequences have labels overlap greater than a threshold, we consider them to be in the similar batch during contrastive learning.

##### Algorithm 1

Algorithm 1 Multi-Label Supervised Pair Creation for RNA Sequences

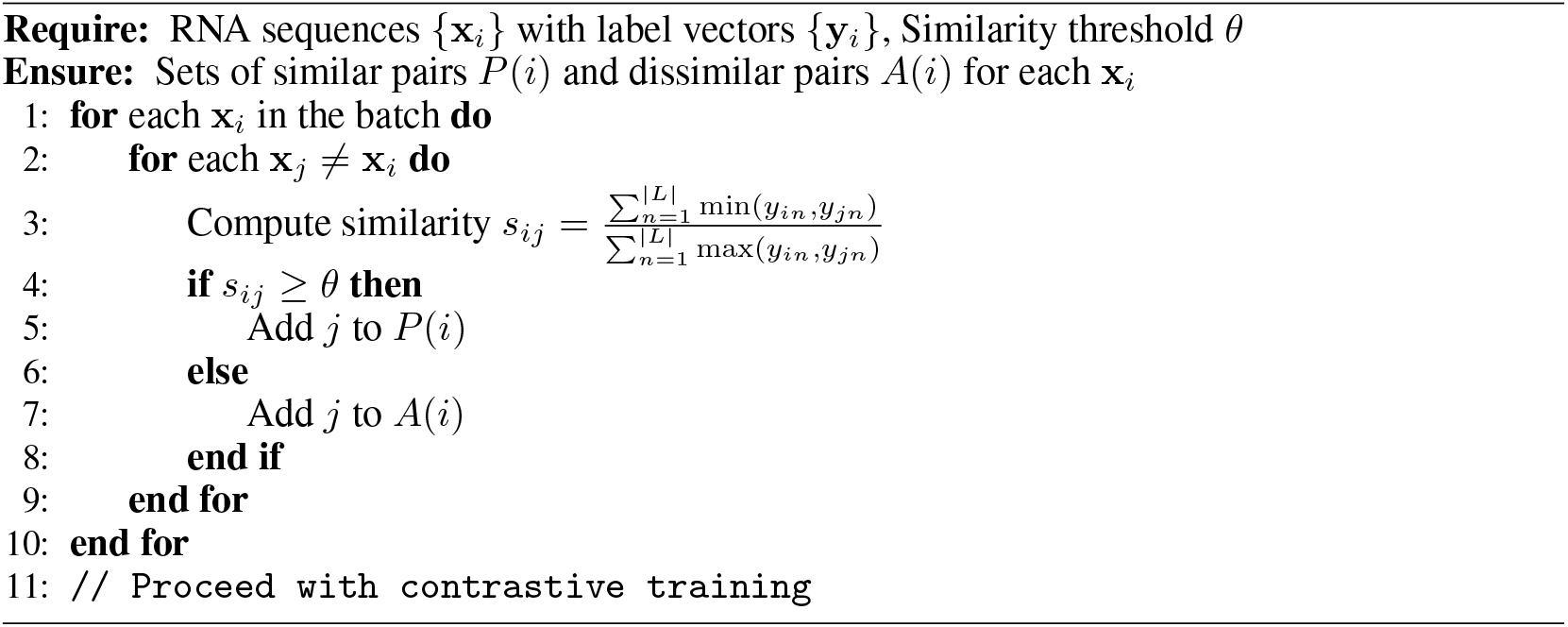

#### Prediction Head with ML-Decoder

We use ML-Decoder [12] for the classification head instead of general classifiers, as it is designed for multi-label classification, replacing the self-attention in traditional transformer-decoders. It takes the output embeddings **E** from a fine-tuned encoder (trained with supervised contrastive loss) and uses group queries **Q**_*k*_ to interact with these embeddings, producing logits **L**_*i*_ for each class.

### 2.4 Complete Training Pipeline of LOCAS

The training of LOCAS involves two steps (Figure 1): **First**, the encoder is fine-tuned using supervised contrastive training to learn embeddings where RNA sequences with similar labels are closer together. **Second**, the encoder’s outputs are frozen and fed into the ML-Decoder, which uses cross-attention to predict label probabilities. The predictions are then compared with ground truth labels to compute a loss, refining the model’s accuracy.

**Figure 1.**
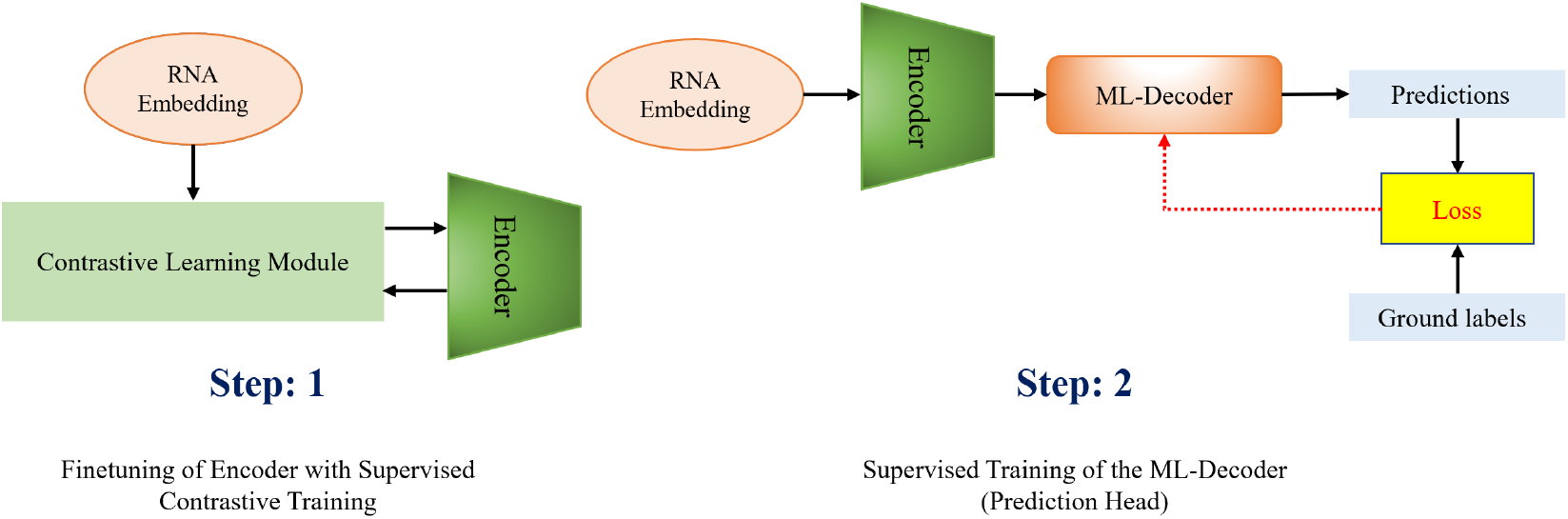
Overall training pipeline for LOCUS. In the first step, the encoder is finetuned with supervised contrastive learning. In the second step the encoder’s output is fed into the ML-decoder for final prediction.

## 3 Results

For RNA subcellular localization prediction, key metrics include Example-Based Accuracy, Average Precision, Coverage, One-Error, Ranking Loss, and Hamming Loss. These metrics evaluate model performance in correctly identifying, ranking, and covering multiple subcellular compartments associated with each RNA sequence.

### Comparison with State of the Art Approaches

The comparison in Figure 2 shows that LOCAS outperforms state-of-the-art models like DM3Loc, Allocator, and Clarion in RNA subcellular localization across several metrics. LOCAS achieves the highest accuracy (Acc_Exam of 0.75) and average precision (0.9434), indicating superior ability in correctly identifying RNA locations. It also records the lowest coverage (2.1262), one-error rate (0.0092), ranking loss (0.0804), and hamming loss (0.1401), reflecting fewer errors and better ranking of correct labels. These results demonstrate that LOCAS provides a more accurate, reliable, and efficient classification of RNA sequences than existing methods. A detailed class-wise performance analysis follows in the next section.

**Figure 2.**
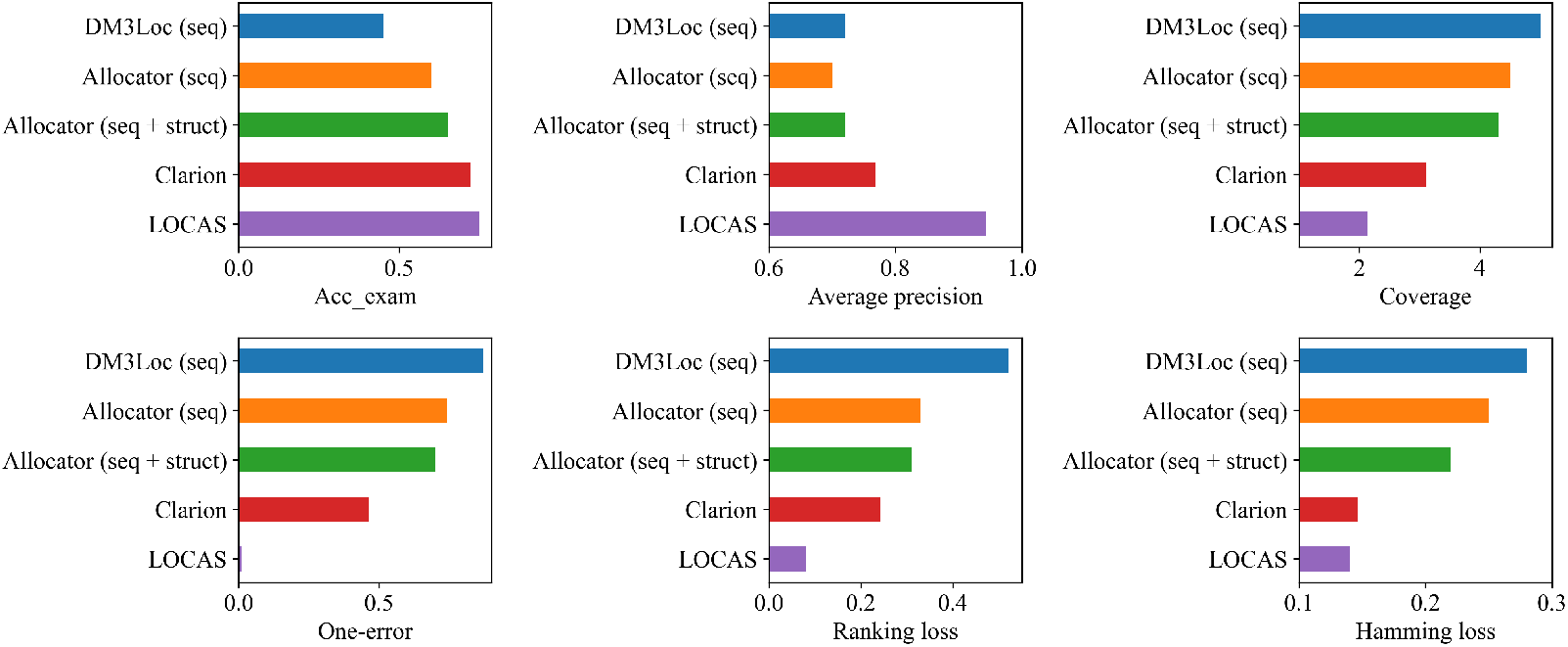
Performance comparison of LOCAS with previously proposed methods for mRNA sub-celuller localization prediction on different metrics.

### Performance on Individual Classes

This section presents a class-wise performance analysis of LOCAS using the Matthews Correlation Coefficient (MCC) in Table 1. LOCAS demonstrates a clear advantage in most RNA localization tasks, such as predicting nuclear RNA, where it achieves an MCC of 0.605, outperforming other methods like DM3Loc and RNATracker. For the exosome class, LOCAS excels with an MCC of 0.501, while other methods fail to make meaningful predictions. LOCAS also shows strong performance in cytosol and ribosome localizations, with MCC values of 0.597 and 0.421, respectively. However, for the ER class, all the methods performs significantly poor due to the scarcity of positive samples, which limits the effectiveness of the supervised contrastive loss in distinguishing this class. Even though iLoc-mRNA[21] performs better for the ER class, performance on the other classes suggest that iLoc-mRNA is highly biased to classify only the ER class better.

**Table 1.** Comparison of MCC values across different methods.

| Method | Nucleus | Exosome | Cytosol | Ribosome | Membrane | ER |
| --- | --- | --- | --- | --- | --- | --- |
| DM3Loc [15] | 0.386 | 0.074 | 0.287 | 0.355 | 0.312 | 0.205 |
| RNATracker [18] | 0.345 | 0.000 | 0.138 | 0.270 | 0.193 | 0.000 |
| mRNALoc [4] | 0.150 | 0.000 | -0.029 | - | - | -0.148 |
| iLoc-mRNA [21] | 0.052 | - | 0.025 | 0.390 | - | **0.376** |
| **LOCAS** | **0.605** | **0.501** | **0.597** | **0.421** | **0.468** | 0.000 |

### 3.1 Ablation Study

To evaluate the contributions of different components in LOCAS, we conducted an ablation study, focusing first on the impact of the supervised contrastive learning (SCL) approach. Results in Table 2, show that omitting SCL leads to a significant drop in performance across all metrics, with MCC scores falling to zero for some classes. For instance, without SCL, the MCC score for the nucleus class drops to 0.142, below that of most other methods, demonstrating the importance of SCL in capturing the contextual information of RNA sequences.

**Table 2.** Comparison of MCC scores for different RNA subcellular localization classes with and without Supervised Contrastive Learning (SCL).

| Method | Nucleus | Exosome | Cytosol | Ribosome | Membrane | End. Reticulum |
| --- | --- | --- | --- | --- | --- | --- |
| **Without SCL** | 0.142 | 0.000 | 0.130 | 0.005 | 0.000 | 0.000 |
| **With SCL** | 0.605 | 0.501 | 0.597 | 0.421 | 0.468 | 0.000 |

We also examined the effect of varying the overlapping threshold in clustering. As shown in Figure Table 3, different thresholds yield varying numbers of clusters and distribution patterns. For a threshold of 0.3 or 0.5, the clustering is heavily biased towards one cluster, while at 0.8, there are 46 clusters with a more balanced, long-tailed distribution. Table 3 confirms that a threshold of 0.8 achieves the best downstream performance, indicating the importance of selecting an appropriate threshold for effective clustering and performance.

**Table 3.**
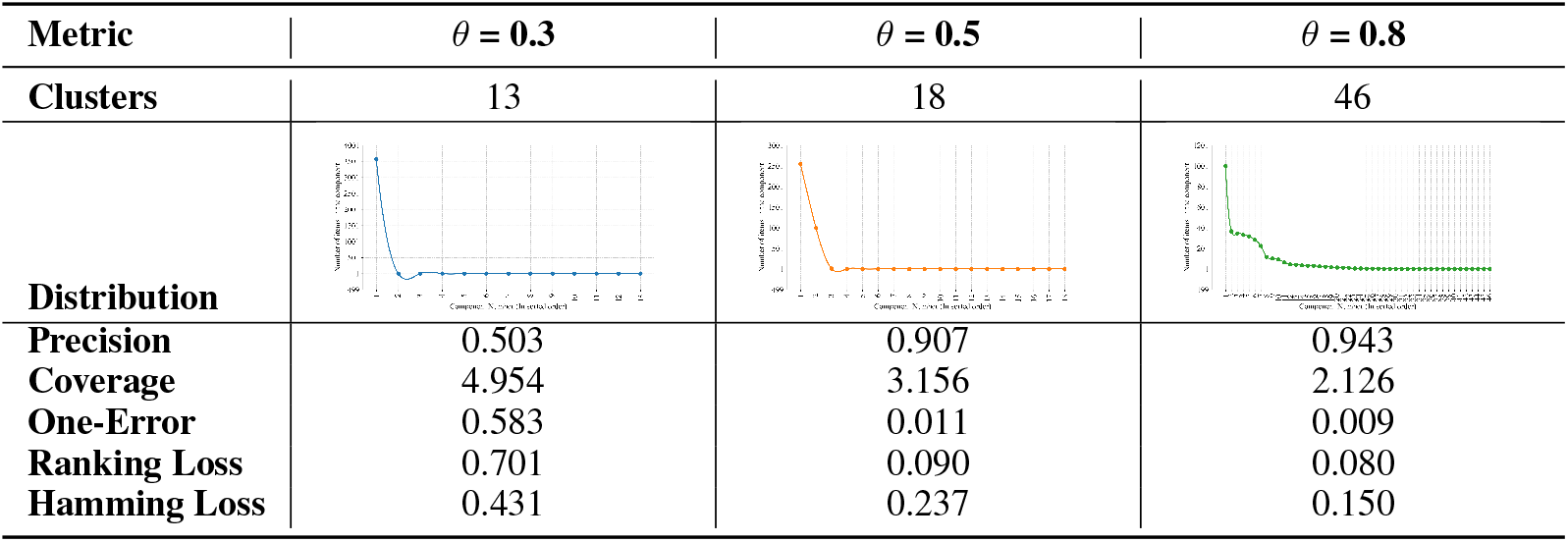
Comparison of Clustering and Performance Metrics for Different threshold Values (*θ*)

| Metric | $\theta = 0.3$ | $\theta = 0.5$ | $\theta = 0.8$ |
| --- | --- | --- | --- |
| **Clusters** | 13 | 18 | 46 |
| **Distribution** |  |  |  |
| **Precision** | 0.503 | 0.907 | 0.943 |
| **Coverage** | 4.954 | 3.156 | 2.126 |
| **One-Error** | 0.583 | 0.011 | 0.009 |
| **Ranking Loss** | 0.701 | 0.090 | 0.080 |
| **Hamming Loss** | 0.431 | 0.237 | 0.150 |

## 4 Conclusion and Future Work

In this work, we present LOCAS, a novel framework for RNA subcellular localization that integrates an RNA language model, supervised contrastive learning (SCL), and a specialized multi-label classification head to achieve state-of-the-art results. LOCAS effectively captures complex relationships in RNA sequences, addressing challenges in multi-label classification and natural label overlap.

Despite its success, LOCAS has certain limitations. The batch-wise handling in SCL may be ineffective when batches lack sufficient sequences from the same class, potentially hindering the model’s ability to learn robust representations. Additionally, the current approach does not consider RNA’s 3D structural information, which could further refine localization predictions.

Future research could focus on developing improved batching strategies to ensure more consistent data distribution and incorporating 3D structural data to complement the sequential features, potentially enhancing both accuracy and generalizability in RNA localization tasks.

## References

[1] Y. Bi, F. Li, X. Guo, Z. Wang, T. Pan, Y. Guo, G. I. Webb, J. Yao, C. Jia, and J. Song. Clarion is a multi-label problem transformation method for identifying mrna subcellular localizations. Briefings in Bioinformatics, 23, 2022.

[2] Son D Dao, Ethan Zhao, Dinh Phung, and Jianfei Cai. Multi-label image classification with contrastive learning. arXiv preprint arXiv:2107.11626, 2021.

[3] CM Di Liegro, G Schiera, and I Di Liegro. Regulation of mrna transport, localization and translation in the nervous system of mammals. International Journal of Molecular Medicine, 33:747–762, 2014.

[4] A Garg, N Singhal, R Kumar, and M Kumar. mrnaloc: a novel machine-learning based in-silico tool to predict mrna subcellular localization. Nucleic Acids Research, 48:W239–W243, 2020.

[5] WR Jeffery, CR Tomlinson, and RD Brodeur. Localization of actin messenger rna during early ascidian development. Developmental Biology, 99:408–417, 1983.

[6] Fuyi Li, Yue Bi, Xudong Guo, Xiaolan Tan, Cong Wang, and Shirui Pan. Advancing mRNA subcellular localization prediction with graph neural network and RNA structure. Bioinformatics, 40(8):btae504, 08 2024.

[7] J Li, L Zhang, S He, F Guo, and Q Zou. Sublocep: a novel ensemble predictor of subcellular localization of eukaryotic mrna based on machine learning. Briefings in Bioinformatics, 22:bbaa401, 2021.

[8] F MacRitchie. Physicochemical properties of wheat proteins in relation to functionality. In Advances in food and nutrition research, volume 36, pages 1–87. Elsevier, 1992.

[9] LA Mingle, NN Okuhama, J Shi, RH Singer, J Condeelis, and G Liu. Localization of all seven messenger rnas for the actin-polymerization nucleator arp2/3 complex in the protrusions of fibroblasts. Journal of Cell Science, 118:2425–2433, 2005.

[10] Jonathan Mojoo and Takio Kurita. Deep metric learning for multi-label and multi-object image retrieval. IEICE TRANSACTIONS on Information and Systems, 104(6):873–880, 2021.

[11] Rafael Josip Penić, Tin Vlašić, Roland G Huber, Yue Wan, and Mile Šikić. Rinalmo: General-purpose rna language models can generalize well on structure prediction tasks. arXiv preprint arXiv:2403.00043, 2024.

[12] Tal Ridnik, Gilad Sharir, Avi Ben-Cohen, Emanuel Ben-Baruch, and Asaf Noy. Ml-decoder: Scalable and versatile classification head. In Proceedings of the IEEE/CVF winter conference on applications of computer vision, pages 32–41, 2023.

[13] Md Toki Tahmid, Haz Sameen Shahgir, Sazan Mahbub, Yue Dong, and Md Shamsuzzoha Bayzid. Birna-bert allows efficient rna language modeling with adaptive tokenization. bioRxiv, pages 2024–07, 2024.

[14] Chao Wang and Quan Zou. Prediction of protein solubility based on sequence physicochemical patterns and distributed representation information with deepsolue. BMC biology, 21(1):12, 2023.

[15] D. Wang, Z. Zhang, Y. Jiang, Z. Mao, D. Wang, H. Lin, and D. Xu. Dm3loc: multi-label mrna subcellular localization prediction and analysis based on multi-head self-attention mechanism. Nucleic Acids Research, 49:e46, 2021.

[16] Jiawei Wang, Bingjiao Yang, Jerico Revote, Andre Leier, Tatiana T Marquez-Lago, Geoffrey Webb, Jiangning Song, Kuo-Chen Chou, and Trevor Lithgow. Possum: a bioinformatics toolkit for generating numerical sequence feature descriptors based on pssm profiles. Bioinformatics, 33(17):2756–2758, 2017.

[17] Z Yan, E Lecuyer, and M Blanchette. Prediction of mrna subcellular localization using deep recurrent neural networks. Bioinformatics, 35:i333–i342, 2019.

[18] Zichao Yan, Eric Lécuyer, and Mathieu Blanchette. Prediction of mrna subcellular localization using deep recurrent neural networks. Bioinformatics, 35(14):i333–i342, 2019.

[19] T. Zhang, P. Tan, L. Wang, N. Jin, Y. Li, L. Zhang, H. Yang, Z. Hu, L. Zhang, C. Hu, C. Li, K. Qian, C. Zhang, Y. Huang, K. Li, H. Lin, and D. Wang. Rnalocate: a resource for rna subcellular localizations. Nucleic Acids Research, 45:D135–D138, 2016.

[20] T Zhang, P Tan, L Wang, N Jin, Y Li, L Zhang, H Yang, Z Hu, L Zhang, C Hu, C Li, K Qian, C Zhang, Y Huang, K Li, H Lin, and D Wang. Rnalocate: a resource for rna subcellular localizations. Nucleic Acids Research, 45:D135–D138, 2017.

[21] Zhao-Yue Zhang, Yu-He Yang, Hui Ding, Dong Wang, Wei Chen, and Hao Lin. Design powerful predictor for mrna subcellular location prediction in homo sapiens. Briefings in Bioinformatics, 22(1):526–535, 2021.

[22] ZY Zhang, YH Yang, H Ding, D Wang, W Chen, and H Lin. Design powerful predictor for mrna subcellular location prediction in homo sapiens. Briefings in Bioinformatics, 22:526–535, 2021.

